# Sex and timing of gonadectomy relative to puberty interact to influence weight, body composition, and feeding behaviors in mice

**DOI:** 10.1101/2022.12.13.520312

**Authors:** Courtney M. Klappenbach, Qing Wang, Allison L. Jensen, Nicholas C. Glodosky, Kristen Delevich

## Abstract

Gonadal sex steroids are important regulators of energy balance in adult rodents, and gonadectomy (GDX) has opposing effects on weight gain in sexually mature males and females. Puberty is associated with the emergence of sex differences in weight, body composition, and feeding behaviors, yet the role of gonadal hormones at puberty remains unclear. To address this, we performed GDX or sham surgery in male and female C57Bl/6 mice at postnatal day (P)25 (prepubertal) or P60 (postpubertal) timepoints and measured weight and body composition for 35 days, after which *ad libitum* and operant food intake was measured using Feeding Experimentation Device 3 (FED3s) in the home cage. Consistent with previous studies, postpubertal GDX caused weight gain in females and weight loss in males and increased adiposity in both sexes. However, prepubertal GDX decreased weight gain and altered body composition across the adolescent transition (P25 to P60) in males but had no effect in females. Despite the varied effects on weight, GDX decreased food intake and motivation for food as assessed in operant tasks regardless of sex or timing of surgery relative to puberty. Our findings indicate that GDX interacts with both sex and age at surgery to influence weight, body composition, and feeding behavior.

**Highlights:**

- Gonadectomy had opposing effects on weight in males and females when performed in adulthood.
- Gonadectomy performed prior to puberty decreased weight gain in males but did not affect weight gain in females.
- Across sex and age at surgery, GDX decreased *ad libitum* food intake and reduced operant responding for food.
- Decreased food intake under effortful conditions was not explained by body weight in GDX mice.

## 1. Introduction

Across species, puberty is associated with rapid somatic growth and changes to body composition (Bitar et al., 2000; Demerath et al., 2004; Sun et al., 2001; Taylor et al., 2010). The onset of puberty involves the reactivation of the hypothalamic pituitary gonadal (HPG) axis which triggers the elevated secretion of gonadal hormones, primarily testosterone (T) from the testes and estradiol (E2) and progesterone from the ovaries (Sisk and Foster, 2004). Gonadal hormones contribute to sex differences in weight and body composition that emerge post puberty (Grunt, 1964; Rogol et al., 2002; Swanson and Van Der Werff Ten Bosch, 1963) as well as sex differences in behaviors that regulate energy balance, including palatable food consumption (Klump et al., 2021; Shomaker et al., 2010) and physical activity (Kennedy and Mitra, 1963; Reiber et al., 2022). Puberty is just one transition that characterizes adolescence, the broader developmental window during which organisms gradually attain adult-like roles and behaviors (Dahl et al., 2018; Piekarski et al., 2017b; Spear, 2000). Adolescence is linked to heightened risk for the onset of eating disorders, which exhibit a significant female sex bias (Klump et al., 2017; Timko et al., 2019) and is increasingly recognized as a period of feeding circuit remodeling (Zeltser, 2018) that may establish persisting trends for energy balance regulation (Ng et al., 2014; Ruiz et al., 2009; Saydah et al., 2013).

Much work has examined the role of gonadal hormones in regulating energy balance in male and female mammals (Mauvais-Jarvis et al., 2013; Wade, 1976). In rodents, surgical removal of the gonads, or GDX, results in significant changes to weight and body composition, mediated by alterations to both intake and expenditure sides of the energy balance equation (Wade, 1976). Existing literature has primarily focused on the effects of GDX in sexually mature rodents, which has opposing effects on weight and food intake in males and females (Asarian and Geary, 2006; Kakolewski et al., 1968). In females, ovariectomy (OVX) and the subsequent reduction in circulating estrogens increases weight gain and fat accumulation in a manner that is dependent on estrogen receptor *α* (ERα) signaling in the brain (Harris et al., 2002; Mamounis et al., 2014; Roepke, 2009; Santollo et al., 2007; Wegorzewska et al., 2008). In male rodents, orchiectomy (ORX) and the reduction in circulating androgens lead to weight loss and increased fat accumulation (Dubois et al., 2016; Gentry and Wade, 1976). The effects of ORX on energy balance appear to be mediated by both androgenic and estrogenic mechanisms, via the aromatization of T to E2 (Jardí et al., 2018a).

Compared to adulthood and the perinatal period, less is known about the influence of gonadal hormones during peripuberty on energy balance. Decades ago it was reported that prepubertal female rats appear resistant to the anorexic effects of estradiol (Wade and Zucker, 1970), but the mechanisms remain unclear (Wade, 1974; Zucker, 1972). Subsequent research, primarily pertaining to reproductive, social, and anxiety-related behaviors, demonstrated that adolescence is a second organizational window during which gonadal sex steroids may exert long-lasting effects and shape how the brain responds to sex steroid hormones in adulthood (Delevich and Wilbrecht, 2020; Schulz and Sisk, 2016). With respect to energy balance, this raises the question of whether the refractoriness of the prepubertal female brain to exogenous sex steroid hormones could be due to a lack of peripubertal organization. A first step to assess potential organizational effects at puberty is to compare the effects of GDX performed before vs. after the peripubertal window (Sisk and Romeo, 2019). To date, there is little data directly comparing the effects of pre-vs. postpubertal GDX in male and female mice. Delineating sex-dependent influences of gonadal hormones at puberty on energy balance will be important for understanding the causal factors that contribute to sex differences in metabolic disease and psychiatric eating disorders.

In addition to changes in growth and metabolism, peripuberty may be a sensitive period for the organization of adult behaviors across multiple domains including social, cognitive, and affective (Delevich and Wilbrecht, 2020; Larsen and Luna, 2018; Piekarski et al., 2017b; Schulz and Sisk, 2016). In particular, there is a growing interest in the influence of puberty and gonadal sex steroids on cognitive and decision-making processes (Delevich et al., 2021; Laube et al., 2017; Master et al., 2020; Orsini et al., 2022; Piekarski et al., 2017a). As food is a common primary reinforcer, it is important to understand how gonadal sex steroids influence motivation or willingness to work for food within contexts relevant to operant decision-making tasks. Therefore, in addition to *ad libitum* food intake, we assessed operant responding (nose poking) for food under increasing schedules of reinforcement to assess the influence of sex, gonadal status, and age on motivation to work for food.

Finally, to-date, most studies that have examined how gonadal hormones influence detailed patterns of food intake and/or operant responding for food have been performed in rats. Species differences in feeding behaviors, as well puberty (Bell, 2018) and estrous-cycle dependent regulation of energy homeostasis suggest that findings in rats do not simply translate to mice (Santollo and Daniels, 2015; Witte et al., 2010). Given the abundance of genetic tools available in mice, detailed characterization of the influence of sex, gonadal status, and age on energy balance are necessary to inform future mechanistic studies.

Here, we compared the effect of prepubertal vs. postpubertal GDX on weight gain, body composition, food intake, and operant responding for food in both male and female C57Bl/6 mice. We performed GDX at pre- or postpubertal ages and compared behavior after a consistent “gonad free” duration of 35 days post-surgery (dps). These experiments enabled us to directly examine how sex and age at surgery influenced the impact of GDX on weight gain, body composition, and feeding-related behaviors in mice.

## 2. Methods

### 2.1 Subjects

Male and female C57BL/6 mice (Charles River) were bred in-house. All mice were weaned on postnatal day (P)21 and housed in groups of 2-3 same-sex siblings on a 12:12 h reversed light:dark cycle (lights on at 1900 h) in standard ventilated polycarbonate cages with Biofresh cellulose bedding. Mice were provided *ad libitum* access to standard chow (Purina 5001) that contains ~5% fat and water in the home cage until pellet-based food intake (FI) testing which provided 5-TUM grain-based enrichment pellets (TestDiet) that contains 3.8% fat. Water was available *ad libitum* during pellet-based FI studies. All procedures were approved by the Washington State University Institutional Animal Care and Use Committee and conformed to principles outlined by the NIH Guide for the Care and Use of Laboratory Animals.

### 2.2 Experimental design

At P25 or P60, male and female mice were randomly assigned to undergo GDX or sham surgery (details below). Mice were weighed daily for one week following surgery, and then every other day until FI was measured from P60–7 or P95–102, respectively. Body composition measurements (baseline, midpoint, and endpoint) were taken using a Minispec LF (Bruker, Billerica, MA, USA). Mice were euthanized at the end of the experiment, at which point serum was collected and the uterus or seminal vesicles were dissected and weighed. An experimental timeline detailing the procedure is provided in Fig. 1. Subjects in this study included 19 P25 sham male, 15 P25 ORX male, 9 P60 sham male, 10 P60 ORX male, 11 P25 sham female, 11 P25 OVX female, 12 P60 sham female, and 12 P60 OVX female mice, for a total of 99 mice. For a separate cohort of female mice, body weight was measured from P25 to P95 and then mice were euthanized. This cohort included 8 P25 sham and 8 P25 OVX mice. One of these OVX mice was removed from analyses after being identified as an outlier in the uterus weight analysis (ROUT: Q = 1%). Some measurements include fewer mice per group, see figure legends for exact numbers per group.

**Figure 1.**
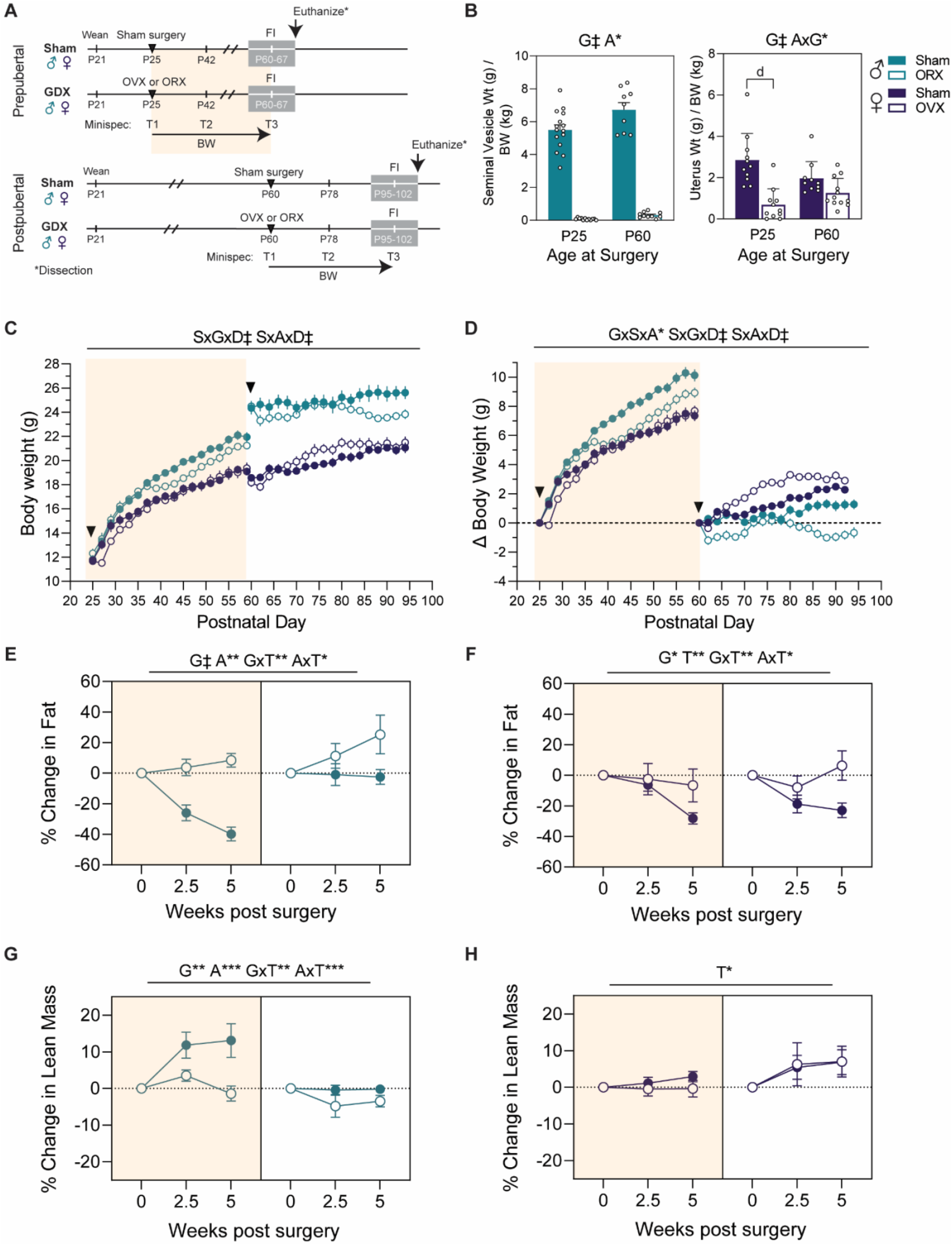
Sex-dependent effects of age at GDX on weight and body composition. **A:** Experimental design. For prepubertal age groups, sham or GDX surgery was performed at P25. For the postpubertal groups, surgery was performed at P60. Food intake (FI) was measured for 6 days at 35 dps (grey box). Body composition was measured just prior to surgery (T1), 2.5 weeks (T2), and 5 weeks post-surgery (T3). From 0 – 35 dps BW was measured. **B:** Weight in g of reproductive organs (seminal vesicle for males, uterus for females) normalized to BW in kg on the day of collection. Males: n = 9-15 mice / group, females: n = 10-12 mice / group. **C:** Raw BW in prepubertal (yellow shading) and postpubertal surgery groups. **D:** Δ BW in prepubertal vs. postpubertal surgery groups. For panels CD, n = 9-17 mice / group and arrows indicate day of surgery. **E:** Post-surgery % fat change in males. **F:** Post-surgery % fat change in females. **G:** Post-surgery % lean mass change in males. **H:** Post-surgery % lean mass change in females. For panels E and G (males), n = 9-12 mice / group. For panels F and H (females), n = 9-12 mice / group. Statistics above panels B and E-H represent three-way ANOVAs for gonadal status (G), sex (S), and age at surgery (A). Statistics above panels C-D represents three-way interactions from linear mixed models with fixed effects of dps (D), age at surgery (A), sex (S), and gonadal status (G) and a random effect for the repeated measure of mouse. For all panels: * *p* < .05, ** *p* < .01, *** *p* < .001, ‡ *p*< .0001 and bar/points represent mean ± SEM.

### 2.3 Gonadectomy

To compare the effects of GDX prior to puberty onset versus after puberty, we performed sham or GDX surgeries at P25 and P60. In mice, vaginal opening is the earliest outward marker of puberty that occurs when estradiol causes the dissolution of a layer of cells that covers the vagina. Prior to P25 ovariectomy (OVX), all female mice were visually inspected to confirm that they had not undergone vaginal opening. In addition, we performed histological analysis of ovaries collected from a separate group of P25 female C57Bl/6 mice and did not observe a corpus luteum in any of the samples (data not shown). In males, the first outward sign of puberty appears later than in females, with the separation of the prepupice from the glans penis at approximately P36 (Deboer and Li, 2011). In a subset of P25 males who received orchiectomies (ORX), we collected samples from the cauda epididymis and visually inspected them under a microscope. While we observed immature spermatids we did not observe mature sperm at this age.

Prior to surgery, mice were injected with 0.05 mg/kg buprenorphine HCl and 10 mg/kg meloxicam subcutaneously and were anesthetized with 1-2% isoflurane during surgery. The incision area was shaved and scrubbed with ethanol and betadine. Ophthalmic ointment was placed over the eyes to prevent drying. A 1 cm incision was made with a scalpel in the lower abdomen across the midline to access the abdominal cavity. The ovaries or testes were clamped off from the uterine horn or spermatic cord, respectively, with locking forceps and ligated with sterile sutures. After ligation, the gonads were excised with a scalpel. The muscle and skin layers were sutured, and wound clips were placed over the incision for 7-10 days to allow the incision to heal. Additional injections of 0.05 mg/kg buprenorphine HCl and 10 mg/kg meloxicam was given 24 and 48h after surgery. Sham control surgeries were identical to gonadectomies except that the gonads were simply visualized and were not clamped, ligated, or excised. Mice were allowed to recover on a heating pad until ambulatory and were post-surgically monitored for 7-10 days to check for normal weight gain and signs of discomfort/distress. Mice were co-housed with 1-2 same-sex siblings who received the same surgical treatment.

### 2.4 Reproductive organ dissection

At the end of the experiments, mice were euthanized using CO2 after which a cardiac puncture was performed to collect blood and serum was prepared and frozen at −80C. To verify that OVX significantly reduced circulating levels of estradiol, uteri from sham and OVX mice were dissected and weighed. Estradiol is known to influence uterine weight through an ERα-mediated mechanism (Kurita et al., 2001). To verify that ORX significantly reduce circulating levels of testosterone, seminal vesicles were dissected and weighed. In some cases, the seminal vesicles of ORX males could not be located, in which case 0.0 g was recorded.

### 2.5 Food intake monitoring

Feeding Experimentation Devices (FED3) (Matikainen-Ankney, 2021) were purchased from Open Ephys (Atlanta, Georgia USA). Devices were programmed using the Arduino interface and FED3 library code (FreeFeeding, FixedRatio1, and ClosedEconomy_PR1). Mice were moved into single-housed cages with a FED3 35 days after surgery. Iso-bedding was used in place of normal bedding to avoid blockage of FED3 dispensers. Mice were checked daily for the duration of the experiment. Daily checks included checking the FED3 for potential jams, weighing the mouse, weighing the FED3, and refilling pellets as needed. In the case of device jamming, 1-2 days of FED data were dropped from the study to remove the period of time when the device was jammed. In some instances, mice were removed from the study after jamming due to weight loss, and only data prior to the time of device jamming is included.

FED3 testing began with the devices on free-feeding mode for three days (experimental days 1-3). In free-feeding mode, FED3 dispenses a pellet each time one is removed. The mode was then updated to fixed ratio 1 (FR1) for two days (experimental days 4 and 5). In FR1 mode, a pellet is dispensed every time there is a left nose poke by the mouse. Devices were then switched to closed economy mode for one day (experimental day 6). Closed economy mode is a form of progressive ratio testing to evaluate food motivation. The number of left pokes required to dispense a pellet (fixed ratio) increases linearly from 1. After 30 minutes of inactivity (no nose pokes), the ratio resets to 1. An auditory and visual stimulus (200 ms tone and light) is given by the device when a correct poke is made in FR1 mode and when the last poke required to dispense a pellet is made in Closed Economy mode. At the beginning of experimental day 4 (when the mode is updated from free-feeding to FR1) the iso-bedding was removed and replaced with fresh bedding.

### 2.6 Data analysis and statistics

Csv files were collected from FED3 and processed using the FED3 Viz software (Matikainen-Ankney, 2021), excel, and in-house R Studio scripts. For FR1 performance curves examining pokes made vs percent correct, data was binned by every 5 pokes so that the mean percent correct within the 5 poke bin was used for subsequent analysis.

For experiments in which a single data point was collected per animal, groups were compared using a three-way ANOVA with main factors of sex (male or female), gonadal status (sham or GDX), and age at surgery (P25 or P60). We report selected group comparisons to explain interactions of factors when significant. Effect sizes are reported as the standard omega squared ω^2^ which indicates the % of total variation accounted for by the factor or interaction.

For data sets where there were multiple measurements per mouse (repeated measure of mouse) and more than one fixed effect, we used linear mixed models with a random effect of mouse. These models were done in R Studio using the lme4 package. The significance of model effects and interactions were performed with the emmeans package. Model effects and interactions were calculated with the joint_tests() function and *post hoc* comparisons were done with the emmeans() function and specifying pairwise comparisons. The multiple comparison adjustment was done with the rbind() function and the adjustment method ‘sidak’ for Sidak’s multiple comparison test. Reported values for groups represent the mean *±* the standard error of the mean (SEM). Additional statistical information can be found in the supplemental statistical excel file. This includes exact p values, degrees of freedom, and F/t ratios for all ANOVAs / mixed models and *post hoc* comparisons.

## 3. Results

Our study included a 2 x 2 x 2 design, with sex, gonadal status (sham vs. GDX), and age at surgery (P25 vs. P60) as independent factors (Fig. 1A). This study enabled us to examine the influence of testicular and ovarian secretions during different developmental windows. GDX or sham surgery at P60 tested activational effects of gonadal hormones in postpubertal adults, whereas GDX or sham surgery at P25 tested activational and/or organizational effects of gonadal hormones during peri -puberty.

### 3.1 Baseline body weight and endpoint reproductive organ weights

At the end of the experiment, we weighed uteri and seminal vesicles to confirm that OVX and ORX reduced circulating levels of estradiol and testosterone, respectively (Fig. 1B). In males, there was a significant main effect of gonadal status (*p*< .0001, ω^2^ = .91), with ORX males having significantly smaller seminal vesicles than sham males (sham = 5.95 ± 0.28 g / kg, ORX = 0.0002 ± 3.70e-5 g / kg). There was also a main effect of age at surgery, where older animals (P60 surgery age, P95 at collection) had slightly larger seminal vesicles relative to their BW compared to younger animals (P25 surgery age = 3.08 ± 0.56 g / kg, P60 surgery age = 3.35 ± 0.78, *p =* .0105, ω^2^ = .01). In females, there was a significant effect of gonadal status (*p*< .0001, ω^2^ = .36) but also a significant interaction between age at surgery and gonadal status (*p*< .0128, ω^2^ = .09), whereby *post hoc* comparisons were significant for P25 surgery groups (P25 surgery sham = 2.85 ± 0.39 g / kg, P25 surgery OVX = 0.69 ± 0.23 g / kg, *p*< .0001) but not P60 surgery groups (P60 sham surgery = 1.96 ± 0.26 g / kg, P60 OVX surgery = 1.26 ± 0.20 g / kg, *p* = .15).

There was a significant effect of age at surgery (*p*< .0001, ω^2^ = .83) and sex (*p* < .0001, ω^2^ = .10) on baseline BW (Supp. Fig. 1), but no main effect of gonadal status (*p* = .92, ω^2^< .0001). In addition, there was a significant interaction of sex and age at surgery (*p*< .0001, ω^2^ = .07), whereby males and females weighed similarly at P25 (P25 females = 11.71 ± 0.20 g, P25 males = 12.08 ± 0.25 g, *p* = .48), but males weighed more than females at P60 (P60 males = 24.43 ± 0.33 g, P60 females = 18.38 ± 0.21 g, *p* <.0001). These data indicate that age- and sex-matched groups weighed similarly prior to surgery and that sex differences emerge between P25 and P60.

### 3.2 Post-gonadectomy body weight change is dependent on sex and age at surgery

We performed linear mixed effects model for BW (Fig. 1C) and Δ BW (Fig. 1D) post-surgery. Both models indicated significant three-way interactions of days post-surgery (dps) and 1) sex and gonadal status and 2) sex and age at surgery. In addition, Δ BW showed a three-way interaction of sex, age at surgery, and gonadal status. Therefore, follow-up two-way repeated measures ANOVAs were conducted separately for relevant contrasts in male (Supp. Fig. 2A), female (Supp. Fig. 2B), sham (Supp. Fig. 2D), and GDX (Supp. Fig. 2D) groups. For all ANOVAs, there was a significant effect of dps, indicating that BW changed across the 35 day measurement period.

For male mice that had surgery at P25, there was a significant interaction between gonadal status and dps (*p*< .0001, ω^2^ = .01). *Post hoc* comparisons showed that sham mice weighed more than ORX animals from 14 to 26 dps (*p*s < .05) (Supp. Fig. 2A). For male mice that had surgery at P60, there was also a significant interaction of gonadal status and dps (*p*< .0001, ω^2^ = .10). ORX males lost weight shortly following surgery compared to sham males (6 dps: *p* = .0175), recovered, but again weighed less than sham surgery mice from 22-32 dps (*p*s < .05) (Supp. Fig. 2A). For female mice that had surgery at P25, there was no significant effect of gonadal status (*p* = .54, ω^2^ = .002). There was an interaction of gonadal status and dps (*p* = 0.0005, ω^2^ = .01), however the only significant *post hoc* comparison was at 2 dps (*p* = .0264) which likely reflects post-op recovery. By contrast, there was a significant interaction of gonadal status and dps for females who had surgery at P60 (*p*< .0001, ω^2^ = .12) with OVX females gaining significantly more weight than sham, particularly 1-3 weeks post-surgery (10-24 dps: *p*s < .05) (Supp. Fig. 2B). These results show that post but not prepubertal OVX caused significant weight gain. To confirm that this was not due to age differences, we performed prepubertal (P25) OVX or sham surgeries in a separate cohort of mice and measured their BW until P95 (the endpoint of the post-puberty P60 surgery age group). Again, there was no significant effect of gonadal status on weight gain (*p* = .40) (Supp. Fig. 3A).

Next, we assessed sex differences in cumulative weight gain (Δ BW) among mice that underwent sham surgery. For younger mice (P25 surgery age), there was a significant interaction of sex and dps (*p* < .0001, ω^2^ = .02). Males gained more weight than females, starting 12 dps (12-34 dps: *p*s < .01) which corresponds to P37. There was also a significant interaction of sex and dps in the P60 surgery group (*p* < .0001, ω^2^ = .04); females gained slightly more weight than males over the P60–95 period, but no *post hoc* comparisons were significant (Supp. Fig. 2C). These data indicate that there are age-dependent sex differences in weight gain among intact mice, with males gaining significantly more weight than females during adolescence and females gaining more weight than males during young adulthood.

Finally, we assessed sex differences in Δ BW among GDX mice. For mice that underwent prepubertal (P25) GDX there was a significant interaction of sex and dps (*p* = .0046, ω^2^ = .01) (Supp. Fig. 2D). In contrast to the persistent sex difference we observed in sham mice, prepubertal ORX males weighed more than prepubertal OVX females only at early dps (2, 6, 8 dps: *p*s < .05) (Supp. Fig. 2C). As expected from the within sex contrasts, there was a significant interaction of sex and dps in postpubertal (P60) GDX mice (*p* < .0001, ω^2^ = .10), with OVX females gaining significantly more weight than ORX males from 4 dps onward (4-10 dps: *p*s < .01, 12-18 dps: *p*s < .001, 20-32 dps: *p*s < .0001) (Supp. Fig. 2D). These data indicate that prepubertal GDX blocks the sex difference in weight gain during adolescence and that postpubertal GDX has opposing effects on BW in male and female mice.

Finally, we noted that male mice that had sham surgery at P25 weighed less at P60 compared to our P60 group males pre-surgery (P25 male sham weight at P60 = 22.34 ± 0.42, P60 male sham weight at P60 = 24.34 ± 0.53, *p* = .0034) (Supp. Fig 4). This effect was not observed in females and suggests that the growth of male mice was stunted by anesthesia and/or surgery at P25.

### 3.2 Sex-dependent effects of gonadectomy on body composition

To determine the change in fat accumulation and lean mass from pre-surgical baseline over the 5-week period that BW data were collected, we measured body composition just prior to surgery, 2.5 weeks post-surgery (midpoint), and 5 weeks post-surgery (endpoint). Given that we observed sex differences in BW and Δ BW, we analyzed body composition for males and females separately with three-way ANOVAs among gonadal status, age at surgery, and timepoint (5 or 2.5 weeks).

In males, there were significant main effects of gonadal status (*p*< .0001, ω^2^ = .17), age at surgery (*p* = .0017, ω^2^ = .09), but not timepoint (*p* = .62, ω^2^ = .27) on % fat mass (Fig. 1E). In addition, there were significant two-way interactions between age at surgery and timepoint (*p* = .0002, ω^2^ = .05) and between gonadal status and timepoint (*p*< .0001, ω^2^ = .11). Collapsed across age, ORX animals had increased fat mass from baseline whereas sham animals lost or maintained fat (2.5 weeks: sham males = −14.69 ± 5.05%, ORX males = 7.20 ± 4.65%, *p* = .0084; 5 weeks: sham males = −23.00 ± 5.35%, ORX males = 16.09 ± 6.35%, *p*< .0001). Collapsed across gonadal status, adult males (P60 surgery age) increased fat composition compared to adolescent males (P25 surgery age) (2.5 weeks: P25 surgery age males = −10.44 ± 4.83%, P60 surgery age males = 5.51 ± 5.48%, *p* = .10; 5 weeks: P25 surgery age males = −14.66 ± 6.00%, P60 surgery age males = 12.16 ± 7.61%, *p* = .0263).

In females, % change in fat mass showed a significant interaction of gonadal status and timepoint (*p* = .0013, ω^2^ = .06) and age at surgery and timepoint (*p* = .0371, ω^2^ = .03) in addition to main effects of gonadal status (*p* = .0350, ω^2^ = .06) and timepoint (*p* = .0012, ω^2^ = .06). Collapsed across age at surgery, sham females lost fat compared to baseline whereas OVX females maintained fat levels (2.5 weeks: sham females = −13.45 ± 3.91%, OVX females = −5.36 ± 6.13%, *p* = .62; 5 weeks: sham females = −25.18 ± 3.13%, OVX females = 0.14 ± 7.12%, *p* = .0084). These results show that pre- and postpubertal GDX results in greater body fat compared to sham surgery for males and females.

In males, the ANOVA for % change in lean mass indicated significant main effects of age at surgery (*p* = .0008, ω^2^ = .12) and gonadal status (*p* = .0039, ω^2^ = .09), and while there was no main effect of timepoint (*p* = .12, ω^2^ = .02) there were significant two-way interactions between timepoint and age at surgery (*p* = .0004, ω^2^ = .06) and timepoint and gonadal status (*p* = .0026, ω^2^ = .05). Collapsed across surgery age, ORX males gained less lean mass compared to sham males (2.5 weeks: sham males = 6.32 ± 2.43%, ORX males = −0.27 ± 1.80%, *p* = .10; 5 weeks: sham males = 7.10 ± 2.94%, ORX males = −2.32 ± 1.30%, *p* = .0204). In addition, collapsed across gonadal status, adolescent males gained more lean mass compared to adult males (2.5 weeks: P25 surgery age males = 7.49 ± 2.03%, P60 surgery age males = −2.72 ± 1.72%, *p* = .0013; 5 weeks: P25 surgery age males = 5.54 ± 2.83%, P60 surgery age males = −1.91 ± 1.03%, *p* = .06). In females, the ANOVA for % change in lean mass indicated a significant main effect of timepoint only (*p* = .0311, ω^2^ = .03) and no significant interactions. Across all female groups, lean mass generally increased from baseline (2.5 weeks: 3.33 ± 1.91%, 5 weeks: 4.30 ± 1.62%). Therefore, we observed a sex difference in the effect of GDX on lean mass gains, where ORX significantly decreased lean mass gains while OVX did not.

### 3.3 Ad libitum food consumption and patterns

After 35 dps, mice were individually housed and food intake was measured using the Feeding Experimentation Device (FED3) (Matikainen-Ankney et al. 2021), a home-cage automated pellet dispenser that delivers 20 mg grain-based enrichment pellets (TestDiet 5-TUM) (Fig. 2A-B). Standard chow was removed, and mice received their full diet from the FED3. A linear regression of the daily decreases in device weight and weight of recorded pellets delivered had a high goodness of fit (*r*^2^ = .97), indicating that FED3 pellet counts were accurate (Fig. 2C). For the first 3 days, FED3 was set to “free-feeding” (FF) mode during which a pellet was delivered to the magazine each time one was removed. We refer to FF measurements as *ad libitum* food intake.

**Figure 2:**
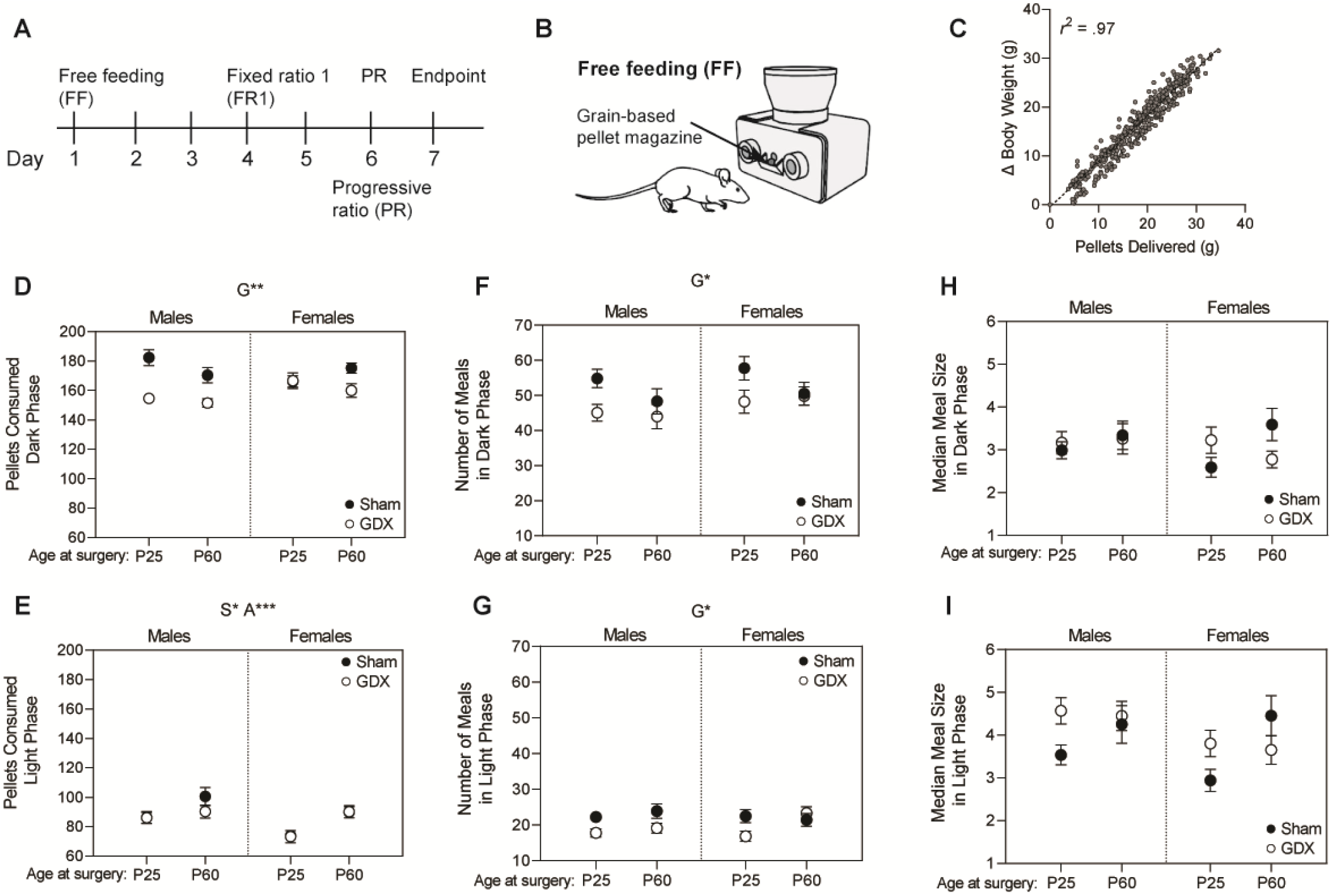
Feeding experimentation devices (FEDs) used for measuring *ad libitum* food consumption. **A:** Timeline of FED modes. FEDs were kept on free-feeding (FF) mode for 3 days, fixed ratio 1 (FR1) mode for 2 days, and finally progressive ratio (PR) testing for 1 day. **B:** Model of mouse and FED in free-feeding (FF) mode. **C:** Correlation between change in device weight and grams of pellets delivered demonstrates that FED devices are accurate. **D:** Pellets consumed from FEDs during the dark phase. **E:** Pellets consumed during the light phase. Meals were defined as pellets consumed within 60 seconds of each other. **F:** Number of meals consumed in the dark phase. **G:** Number of meals consumed in the light phase. **H:** Median size of meals consumed in the dark phase. **I:** Median size of meals in the light phase. Points represent mean ± SEM. Statistics above panels represent linear mixed models with fixed effects of gonadal status (G), sex (S), and age at surgery (A) and a random effect for the repeated measure of mouse (3 days of FF / mouse). For main effects and interactions: * = *p*< .05, ** = *p*< .01, *** = *p*< .001, ‡ = *p*< .0001. For all panels, n = 9-19 mice / group.

To investigate the effects of circadian patterning of our mice groups, we ran a linear mixed model for pellet consumption that included main effects and interactions for the variables of age at surgery, sex, gonadal status, and diurnal cycle (dark or light) as well as a random effect for the repeated measure of mouse (3 days of FF measurements / mouse). This model showed a significant effect of diurnal cycle (*p* < .0001), where mice consumed more pellets in the dark phase compared to the light phase (dark phase = 166.61 ± 1.70, light phase = 85.85 ± 1.54) and several interactions between cycle and other fixed effects including a four-way interaction among sex, age, gonadal status, and cycle (*p* = .0387) (see supplemental statistics file for more details). Therefore, pellet consumption was analyzed separately for the light and dark phases with linear mixed models that included sex, age at surgery, and gonadal status.

For pellets consumed during the dark phase (Fig. 2D), there was a significant effect of gonadal status (*p* = .0019), where sham animals consumed more pellets per dark cycle than GDX animals (sham = 175.00 ± 2.51, GDX = 158.04 ± 2.08). There were no other significant main effects or interactions on dark phase pellet consumption. Interestingly, the number of pellets consumed in the light cycle (Fig. 2E) did not show a significant effect of gonadal status (*p* = .46) but there were significant effects of age at surgery (*p* =.0005) and sex (*p* = .0128). P60 animals (~P95 at test) consumed significantly more pellets than P25 animals (~P60 at test) during the light phase (P25 surgery = 81.11 ± 2.01, P60 surgery = 92.25 ± 2.27), and males consumed more pellets per light phase than females (males = 89.31 ± 2.25, females = 81.85 ± 2.00).

To follow up on these observed cycle differences, we looked at how the proportion of pellets consumed in the dark vs. light phase varied by sex, surgery age, and gonadal status. This was calculated as the number of pellets in the dark phase divided by the total pellets consumed across both phases for each day of FF. Similar to the effects seen in the light phase, there were significant main effects of sex (*p* = .0307) and age at surgery (*p* = .0025) but not gonadal status (*p* = .24) nor any significant interactions. Males and older mice consumed a lower proportion of pellets in the dark phase compared to females (males = .653 ± .0068, females = .673 ± .0063) and younger mice (P60 surgery age = .643 ±.0063, P25 surgery age = .676 ± .0065), respectively. Collectively, these results suggest that GDX selectively decreases pellet consumption during the active (dark) phase, and males and older mice (P95) exhibit attenuated diurnal feeding rhythms compared to females and younger adult mice (P60).

We next examined meal patterning during FF by looking at the number of meals consumed and the size of each meal. Meals were defined as groups of pellets that were consumed within 60 seconds of each other (*i.e*., maximum interpellet interval = 60 s). Since we observed diurnal cycle differences in pellet consumption, we first ran an initial linear mixed model with main effects and interactions of sex, age at surgery, gonadal status, and cycle. For number of meals, we observed a significant effect of cycle (*p*< .0001), where more meals were consumed in the dark than light phase (dark = 50.12 ± 1.09, light = 20.82 ± 0.56) and an interaction between age at surgery and cycle (*p* = .01300). Therefore, we analyzed the number of meals consumed in the light and dark phases separately. In both the light (Fig. 2F) and dark phases (Fig. 2G), linear mixed models showed a significant effect of gonadal status only (dark phase: *p* = .0232, light phase: *p* =.0226), with no other significant effects or interactions. GDX animals consumed fewer meals than sham animals in both phases of the diurnal cycle (dark: sham = 53.40 ± 1.58, GDX = 46.78 ± 1.44; light: sham = 22.40 ± 0.83, GDX = 19.22 ± 0.74).

To investigate meal size, we calculated the median size of all meals consumed per phase (12 hours) so that each mouse had 6 values (3 days of FF x 2 light phases). An initial linear mixed model that included interaction effects of diurnal cycle, sex, gonadal status, and age at surgery, we also observed a main effect of cycle (*p*< .0001). Meals were larger in the light phase compared to the dark phase (dark = 3.10 ± 0.10, light = 3.93 ± 0.12). Due to this main effect, we also analyzed meal size separately for light (Fig. 2H) and dark phases (Fig. 2D), however there were no significant main effects or interactions. Our analysis of meal patterning shows that in both light and dark phases, GDX mice consumed fewer meals of similar size compared to sham mice.

### 3.4 GDX decreases operant responding for food, particularly under high-cost conditions

We hypothesized that GDX-associated decreases in food intake could be due to alterations in the motivation for food. To address this question, we trained mice to operantly respond for food pellets in a fixed-ratio 1 mode (FR1) task (Fig. 3A). One nose poke to the active port triggered pellet delivery, whereas nose pokes to the inactive port were logged but did not result in pellet delivery. An ANOVA for the average total pokes (inactive and active) per day indicated a significant main effect of gonadal status (*p*< .0001, ω^2^ = .17) with GDX mice poking less than sham (sham = 218.35 ± 4.62, GDX = 188.64 ± 5.28) (Fig. 3B).

**Figure 3.**
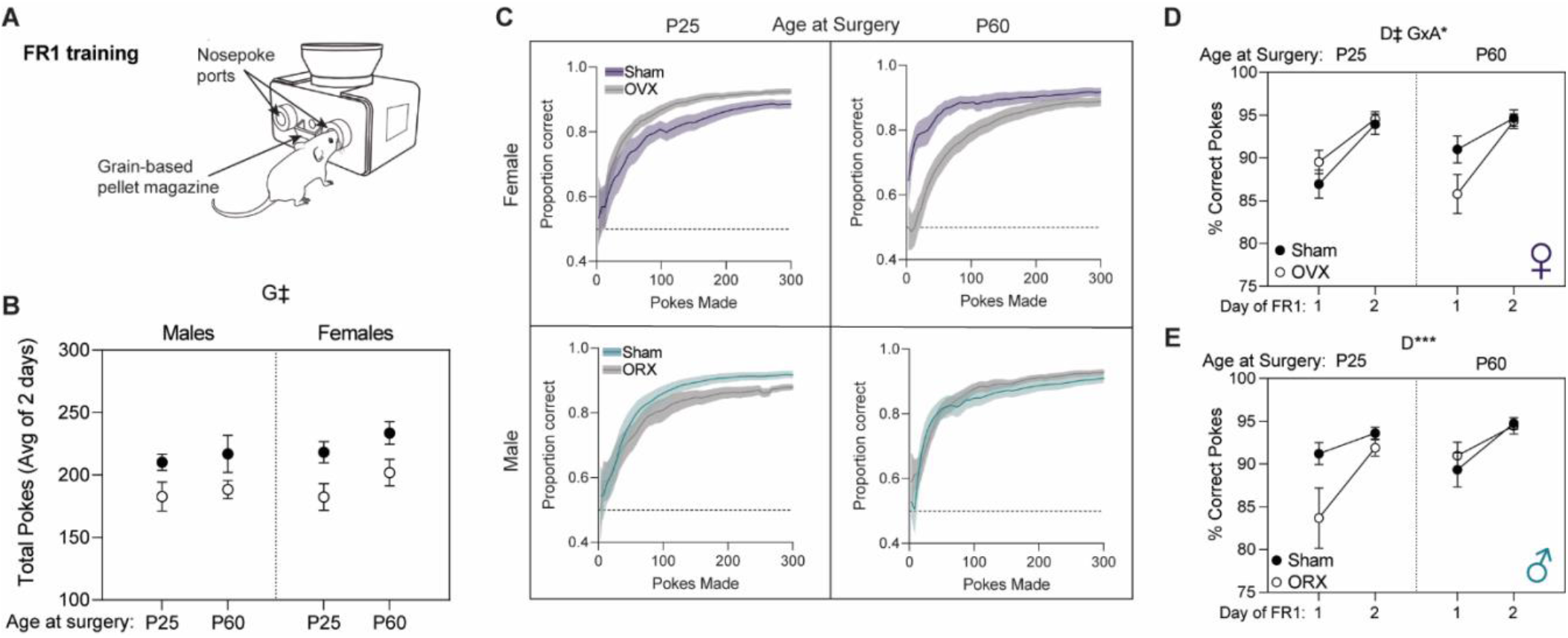
Sex, gonadal status, and age at surgery exert subtle effects on FR1 learning and performance. **A:** Diagram of FR1 training using FED devices. **B:** Average total pokes (active and inactive) per day of two days of FR1 training. **C:** Cumulative proportion correct pokes in FR1 vs total number of pokes made. Data was binned by every 5 pokes and pokes after 300 were trimmed from plot. **D:** % Correct pokes made on days 1 and 2 of FR1 testing in females. **E:** % Correct pokes made on days 1 and 2 of FR1 testing in males. n = 9-17 mice / group. Statistics above panels B, D, and E represent-way ANOVAs between gonadal status (G), sex (S), and age at surgery (A) where * = *p* < .05, *** = *p*< .001, and ‡ = *p* < .0001. Points in figures B, D, and E represent mean ± SEM. Lines in panel C represent mean and error ribbons represent SEM.

To compare FR1 learning we fit cumulative performance curves (proportion correct/cumulative pokes) (Fig. 3C) across groups. We analyzed % correct pokes during FR1 using a three-way ANOVA among task day, gonadal status, and age at surgery. We performed these analyses separately for females (Fig. 3D) and males (Fig. 3E) because of the observed differences in the performance curves (Fig. 3C). In females, prepubertal OVX mice (P25 surgery age) learned FR1 faster (steeper learning curve) than their sham counterparts, whereas the opposite trend was observed in postpubertal OVX mice. This is supported by a significant interaction between gonadal status and age at surgery on % correct pokes (*p* = .0493, ω^2^ = .04), although no sham vs. OVX *post hoc* comparisons were significant (P25 sham = 90.43 ± 1.25%, P25 OVX = 92.02 ± 0.96, p = .59; P60 sham = 92.80 ± 0.98, P60 OVX = 90.03 ± 1.49%, p = .21). There was also a significant main effect of day (*p*< .0001, ω^2^ = .28), where performance improved from day 1 to day 2 (day 1 = 88.18 ± 0.92%, day 2 = 94.33 ± 0.48%). In males, a three-way ANOVA among gonadal status, age at surgery, and day indicated only a significant effect of day (*p* = .0004, ω^2^ = .11; day 1 = 88.61 ± 1.27, day 2 = 93.46 ± 0.44). Overall, these data indicate that all groups rapidly acquired the FR1 task and that interactions between gonadal status and age at surgery impacted FR1 performance in females but not males.

Following 2 days of FR1 training, mice were tested in a progressive ratio (PR) task in which the number of correct pokes required to earn a pellet increased linearly. We used a “closed economy” version of the PR task, in which the reward schedule reset to FR1 after 30 minutes elapsed without interactions (Matikainen-Ankney et al. 2021; Mourra et al. 2020). Analysis of total pokes (active and inactive) during the PR session indicated that there was a main effect of gonadal status (*p*< .0001, ω^2^ = .22) whereby sham mice made more pokes than GDX (sham = 1959.39 ± 88.19, GDX = 1252.79 ± 106.80). There was also a main effect of age at surgery (*p* = .0094, ω^2^ = .05), whereby older mice poked more than younger mice (P25 surgery age = 1463.88 ± 77.78, P60 surgery age = 1809.00 ± 145.42) (Fig. 4A). Examining % correct pokes during PR, there was a significant main effect of gonadal status (*p* = .0014, ω^2^ = .11) with GDX mice exhibiting lower accuracy compared to sham (sham = 96.54 ± 0.27%, GDX = 95.06 ± 0.39%). (Fig. 4B).

**Figure 4.**
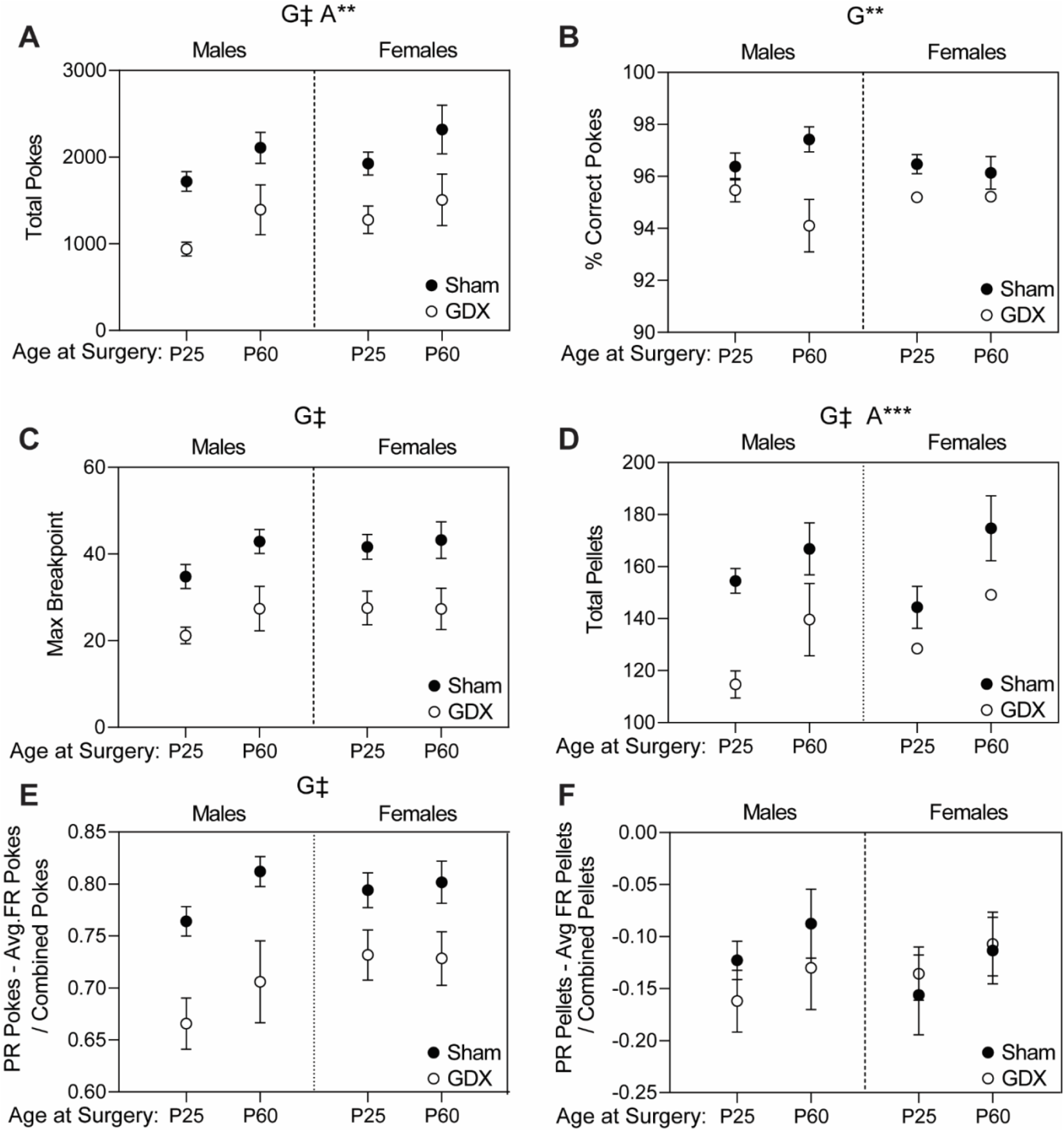
GDX decreases motivation for food in PR task. **A:** Total pokes during 24 hr closed economy PR task. **B:** % correct pokes (accuracy) during PR task. **C:** Maximum breakpoint achieved during PR task. **D:** Total pellets earned during PR task. **E:** Difference in pokes made in FR1 vs PR. Calculated as: pokes made in 1 day of PR – average pokes per day of FR1 divided by the sum of these. **F:** Difference in pellets consumed in FR1 vs PR. Calculated as: pellets consumed in 1 day of PR – average pellets consumed per day of FR1 divided by the sum of these. Data presented as mean ± SEM and were analyzed using three-way ANOVA with gonadal status (G), sex, and age at surgery (A) as factors where ** = *p*< .01, *** = *p* < .001, and ‡ = *p* < .0001. For all panels, n = 9-19 mice / group.

On the closed economy task, mice reach multiple breakpoints when, after 30 min of inactivity, the fixed ratio (pokes required per pellet) resets to 1. We compared the max breakpoint reached in the 24 hours of testing to examine motivation for food. The ANOVA for max breakpoint indicated a significant main effect of gonadal status (*p*< .0001, ω^2^ = .26), with GDX mice exhibiting lower max breakpoints compared to sham (sham = 39.53 ± 1.65, GDX = 25.48 ± 1.91). (Fig. 4C). The ANOVA for total pellets earned during PR indicated significant main effects of gonadal status (*p*< .0001, ω^2^ = .14) and age at surgery (*p* = .0009, ω^2^ = 0.10) (Fig. 4D). Consistent with our results from FF, GDX mice consumed fewer pellets than sham mice (sham = 158.59 ± 4.22, GDX = 131.65 ± 5.06), and older mice earned more pellets during PR than younger mice (P25 surgery age = 136.71 ± 3.77, P60 surgery age = 156.93 ± 6.25).

To examine how food intake was affected by increasing response ratio, we compared daily pellet consumption and nose poking between FR1 and PR conditions. We first calculated the difference in nose pokes between PR (high cost) and FR1 (low cost) as PR nose pokes – FR1 nose pokes / total nose pokes to assess changes in effort. The same calculation was performed for pellet consumption to roughly estimate the elasticity of food demand. As expected, all mice increased responding on PR compared to FR1, as indicated by all values being > 0 (Fig. 4E). However, there was a significant main effect of gonadal status (*p*< .0001, ω^2^ = .21), whereby GDX mice increased their responding during PR to a lesser extent than sham mice (sham = .79 ± .008, GDX = .71 ± .01). By contrast, mice universally reduced food intake on PR compared to FR1, and there was no significant main effect of gonadal status (*p* = .53, ω^2^ =.004) nor any other main effects or interactions. These data indicate that GDX mice exhibited a more thrifty behavioral strategy, expending less effort while maintaining a comparable level of food consumption between FR1 and PR task modes. These findings suggest that regardless of sex or timing relative to puberty, GDX reduces willingness to expend effort for food.

Finally, while all mice lost weight upon switching from FR1 to PR, there was a significant effect of age at surgery on this weight loss (*p* = .0163, ω^2^ = .06) where animals that had surgery at P25 lost more weight than animals that had surgery at P60 (P25 surgery age = −4.06 ± 0.43%, P60 surgery age = −2.41 ± 0.38%) (Supp. Fig. 5).

### 3.5 Changes in food intake relative to BW across increasing cost

Given that sex, gonadal status, and age at surgery influenced BW, we next investigated how food consumption relative to BW varied across groups. For this, we normalized food intake (g) for each day of FED testing to the mouse’s BW at the start of that day. We also wanted to investigate how this measure varied across the three FED modes tested (FF, FR1, PR) as these represent increasing cost / effort required to obtain food. We ran a linear mixed model with main effects and interactions between sex, gonadal status, age at surgery, and task mode (FF, FR1, or PR). This model also had a random effect for the repeated measure of mouse, where each mouse had 6 data points (3 FF, 2 FR1, and 1 PR). In this model, we observed significant main effects of sex (*p*< .0001), gonadal status (*p* = .0008), and mode (*p*< .0001). In addition, there were significant two-way interactions between gonadal status and mode (*p* = .0355) and between age at surgery and mode (*p* =.0056).

The main effect of sex is explained by the observation that females consume more food relative to their body weight than males (females = 0.21 ± 0.003, males = 0.18 ± 0.003) (Fig. 5A). To explore the significant two-way interactions, we plotted these comparisons collapsed across other variables and performed *post hoc* comparisons. For the interaction between gonadal status and mode (Fig. 5B), GDX mice consumed significantly less food relative to their BW than sham mice during FR1 and PR but not FF (FF: sham = 0.23 ± .003, GDX = 0.23 ± .003, *p* = .36; FR1: sham = 0.18 ± .004, GDX = 0.16 ± 0.004, *p* = .0008; PR: sham = 0.14 ± 0.004, GDX = 0.12 ± 0.004, *p* = .0231). This suggests that GDX animals eat less than sham mice under costly conditions (FR1, PR), and that this is not simply explained by BW. To explain the interaction between age at surgery and mode (Fig. 5C), we observed that younger animals (P25 at surgery) consumed more food relative to their BW than older animals (P60 at surgery) in FF, but not FR1 or PR (FF: P25 surgery = 0.235 ± 0.003, P60 surgery = 0.222 ± 0.004, *p* = .0064; FR1: P25 surgery = 0.169 ± 0.004, P60 surgery = 0.166 ± 0.005, *p* = .71; PR: P25 surgery = 0.127 ± 0.004, P60 surgery = 0.134 ± 0.006, *p* = .82). This result indicates that younger animals consumed more food relative to their BW than older animals only under *ad libitum* conditions (FF). Collectively, these results indicate that changes in food intake and motivation observed across sex, gonadal status, and age at surgery are not fully explained by BW.

**Figure 5.**
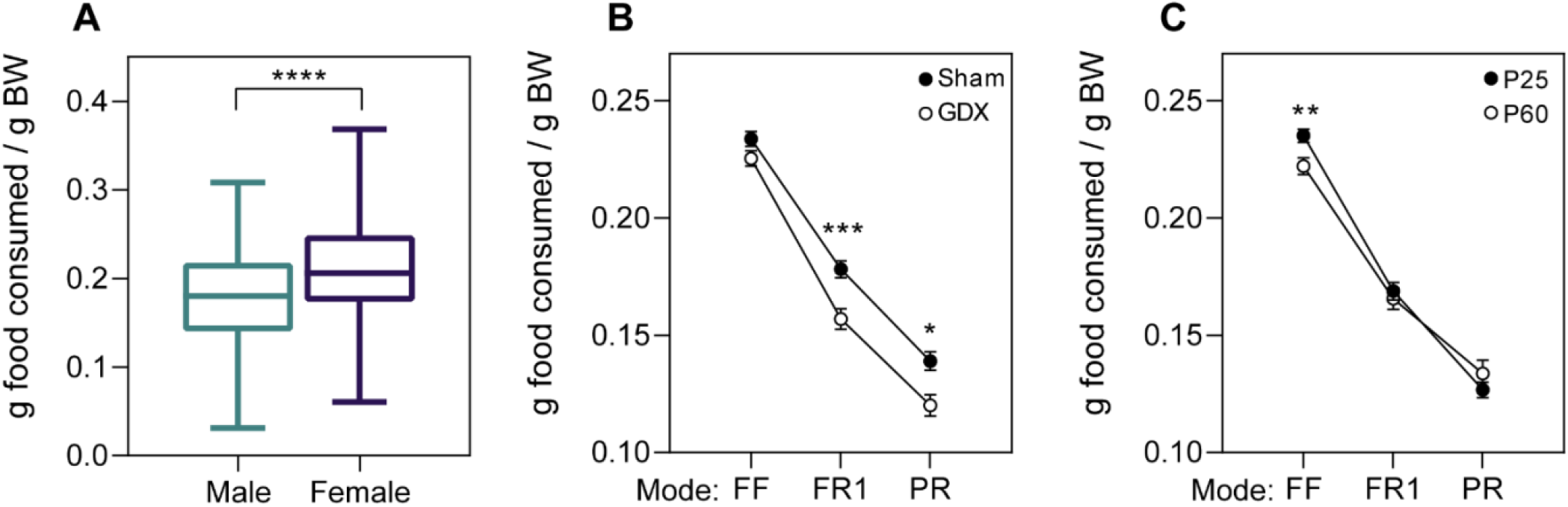
Gonadal status and age at surgery influence food intake relative to body weight differentially across increasing cost. Food intake was normalized to BW for each day of FED testing and a linear mixed model was run with main effects and interactions between sex, age at surgery, gonadal status, and device mode. There was a significant main effect of **A**: sex as well as two significant two-way interactions, **B:** surgery type x mode and **C:** age at surgery x mode. Data presented as mean ± SEM. For *post hoc* comparisons between groups at each mode: * = *p*< .05, ** = *p*< .01, *** = *p*< .001. For main effect of sex: **** = *p*< .0001. For all panels, n = 41-56 mice / group.

## 4. Discussion

Puberty is accompanied by profound morphological, metabolic, and neurobehavioral changes that can have lasting impacts on health and disease. Determining the sex-dependent influence of gonadal hormones during peripuberty will be important for understanding the etiology of sex differences in risk for metabolic disorders, including obesity (Kelly et al., 2008), and eating disorders like anorexia nervosa (Klump, 2013). Here, we compared the effect of pre-vs. postpubertal GDX and sham surgery in male and female mice on physiological outcomes and behavioral regulators of energy balance. Rather than comparing across studies, our experimental design enabled us to draw statistical conclusions regarding the role of sex, GDX, and timing of GDX relative to puberty on multiple aspects of energy balance (Makin and Orban de Xivry, 2019). We found that sex, gonadal status, and age each had independent and, in several cases, interacting effects on measures related to energy balance. To review these findings, we will first discuss influences of sex and age within gonad-intact groups, then discuss how GDX interacts with sex and age to alter weight, body composition, and feeding.

### Sex differences among gonad-intact groups

Comparison of sham surgery (intact) groups revealed sex differences in weight gain and body composition. Males gained significantly more weight than females during the pre-to postpubertal transition (P25 to P60), thus accounting for the emergence of sex differences in BW in young adulthood. Significant sex differences in weight gain were first detected 12 dps (P37), consistent with a previous study in mice (Rathod and Di Fulvio, 2021) and corresponding to a timepoint when plasma testosterone levels are reported to significantly increase in male mice (Wu et al., 2010). Meanwhile, intact females gained more weight compared to males from P60 to P95, consistent with what has previously been described (Moore et al., 2011). In terms of body composition, intact males exhibited the greatest increase in lean mass from P25 to P60 whereas lean mass markedly increased between P60 to P90 in intact females. These results indicate that intact male and female mice exhibit different trajectories of weight gain and body composition changes during the transition from juvenile to mature adult.

In measures of free food intake, males and females consumed a similar number of pellets during the dark phase, consistent with our finding that females consume more food per gram BW. Males consumed proportionally fewer of their daily pellets in the dark phase, suggesting that feeding is more synced to the dark phase in females than males. While we did not examine detailed circadian patterning of food intake, one study found that intact females had an earlier onset and peak of the diurnal feeding rhythm compared to intact males (Chen et al., 2015). Finally, there were no detected sex differences in behavior during PR, indicating that aspects such as accuracy (% correct pokes) and motivation (max breakpoint) are consistent in males and females.

### Age differences among gonad-intact groups

We varied the age at GDX but maintained the duration mice were “gonad free” after which we measured food intake. Therefore, sham groups were assessed either at a young adult (P60) or adult (P95) timepoint, permitting age comparisons within intact mice. As expected, weight gain was significantly greater from P25 to P60 compared to P60 to P95, and there was a main effect of age on several aspects of body composition. In intact males, body fat decreased while lean mass increased from P25 to P60 with little change between P60 to P95. Finally, as previously mentioned, females gained more lean mass from P60 to P95 whereas males predominately gained lean mass from P25 to P60. Collapsed across FED modes, we observed that younger adult mice (P60) consumed more food normalized to BW than older adult (P95) mice. Our “young adult” age of P60 is on the lower age range for a sexually mature adult; given that adolescent mice consume more food per gram BW than adult mice (Moore et al., 2011), we may have detected the tail end of the adolescent period of elevated food intake.

Examining circadian effects on free food intake, the number of pellets consumed during the light phase but not dark phase increased between P60 and P95. These data suggest a shift in diurnal rhythms in food intake between P60 and P95. The underlying mechanism is unclear but could contribute to an increased propensity towards developing obesity with advancing age (Arble et al., 2009). We observed that older mice poked more and consumed more pellets during the PR task than younger adults, but this is likely explained by BW as there was no difference in food intake normalized to BW during PR. One potential confound of interpreting age effects in our study is that mice experienced anesthesia/surgery at different ages, and there is evidence that early postnatal exposure to isoflurane anesthesia can have lasting impacts on behavior (Landin et al., 2019). While the isoflurane concentration cited in this previous study was higher than what we used to achieve surgical anesthesia, we did find that prepubertal sham surgery decreased weight gain relative to unmanipulated males. However, all age effects we report were seen across sex, suggesting that they were not driven by altered growth in the P25 sham surgery males. *GDX effects that are consistent across sex*

In our assays of feeding behaviors, we found that GDX had consistent effects across sex in multiple measures. GDX significantly decreased the number of pellets consumed under *ad libitum* and operant conditions in males and females, regardless of age at surgery. This is consistent with findings in Four Core Genotype mice that GDX reduces food intake, regardless of sex chromosome complement or gonad type (Chen et al., 2015). Given that GDX reduced food intake generally, we compared pellets consumed and nose pokes between PR and FR1 within animal to specifically examine the influence of cost/effort on food intake. Interestingly, there was no significant effect of GDX on elasticity of food demand between PR and FR1 conditions, with GDX and sham mice similarly decreasing their pellet consumption on PR compared to FR1. The effect of GDX primarily manifested as decreased nose poking for food pellets in the face of escalating cost. This was evidenced by reduced max breakpoints during PR and a smaller increase in nose poking when mice were switched from FR1 to PR conditions. These data suggest that GDX shifted both male and female mice towards a less effortful, more thrifty behavioral strategy in pursuit of food. When comparing the effect of escalating cost on pellet consumption vs. nose poking, our data indicate that GDX mice were more economical and expended less energy to maintain relatively stable food intake between FR1 and PR conditions. Importantly, GDX mice reached similar performance in FR1 as sham mice, suggesting that their decreased responding during the PR task was not due to impaired acquisition of the operant response. During PR, however, GDX mice had a lower % of pokes to the active port. While GDX mice maintained a high level of performance (>90%; Fig. 4), this reduced accuracy may reflect decreased engagement or motivation. Indeed, normalized to body weight, GDX mice consumed less food under both “costly” conditions of FR1 and PR compared to sham mice. This effect was not seen under *ad libitum* “free-feeding” conditions where no operant response was required. Despite the different effects that GDX had on weight depending on sex and age at surgery, it caused a clear decrease in the willingness to expend effort for food across all groups examined.

Several studies support our finding of decreased operant responding among OVX females in the PR task. A study by Wang et al. found that estradiol treatment to OVX rats increased perseverative responding in an operant task and promoted a less efficient strategy in pursuit of food reward (Wang et al., 2008). Relatedly, we observed that prepubertal OVX females were less perseverative compared to sham females in an odor-based reversal task that was rewarded by food (Delevich et al., 2021). ERα signaling may negatively regulate cost sensitivity – i.e. enhance food-seeking – as ERα knockout was associated with increased elasticity of food demand in adult female mice (Minervini et al., 2015), but see (Uban et al., 2012). The relationship between estradiol and response persistence may extend to drug rewards and contribute to sex differences in the acquisition and escalation of drug self-administration in (Becker, 2016; Perry et al., 2013). The duration following OVX may affect food motivation, as a study that examined food-seeking behaviors acutely following OVX in adult rats (<14 dps) found that OVX enhanced whereas E2 suppressed lever pressing for sucrose reward (Richard et al., 2017), contrary to our findings.

In a previous study, we observed that prepubertal ORX mice exhibited greatly reduced engagement in an odor-based discrimination task that was rewarded by food, whereas prepubertal OVX produced the opposite effect (Delevich et al., 2020). Our current data help explain reduced task engagement in prepubertal ORX males, but the results from prepubertal OVX females run contrary to what we may have predicted from their performance in an odor-guided foraging task (Delevich et al., 2020). It is possible that decreased anxiety-like behavior we previously observed in prepubertal OVX females accounted for differences in task engagement in the odor-guided foraging task which took place in a large, well-lit arena (Delevich et al., 2020). By testing operant responding for food within the home cage, our current study may have minimized the contribution of anxiety-related behaviors to operant task performance and food seeking behavior.

### Sex-differentiated effects of GDX

Consistent with previous studies, we found that GDX in adult mice had the opposite effect on weight in males and females, with OVX females gaining weight and ORX males losing weight. This difference in weight gain presented despite similar reductions in food intake and increases in adiposity, also consistent with previous observations in mice (Chen et al., 2015). Prepubertal ORX decreased weight gain in males compared to sham, whereas, strikingly, there was no significant difference between prepubertal OVX and sham females. P25 ORX males exhibited a shallower growth curve compared to sham males starting around the age of puberty onset (P37), consistent with what was observed by Petersen (Petersen, 1978). Two existing studies that performed prepubertal GDX in rats reported opposing effects in males and females (Grunt, 1964; Mauvais-Jarvis et al., 2013), with males gaining less weight compared to sham and females gaining more. While we did not observe a significant effect of OVX in prepubertal females, overall, prepubertal GDX eliminated significant sex differences in weight gain that were seen in sham mice. These data suggest that testicular hormones during peripuberty contribute to the emergence of sex differences in weight by early adulthood.

Given the known sex differences in GDX on body composition, we analyzed male and female separately. Generally, ORX caused greater changes in body composition than OVX, greatly increasing body fat and reducing lean mass compared to sham males. In adult males, it has been shown that increased fat mass following ORX can be suppressed by testosterone (T) but not by the non-aromatizable androgen, DHT, suggesting a role of estradiol in reducing fat mass. OVX had subtler effects promoting adiposity and did not significantly affect lean mass.

### Age-dependent effects of GDX

We found that whether GDX was performed before or after puberty determined the effect on weight gain. ORX decreased weight gain over the subsequent 5 weeks whether performed pre- or postpubertally. However, the temporal pattern of this effect differed by age at surgery: prepubertal ORX caused a delayed suppression of weight gain, coinciding with the peripubertal window, whereas postpubertal GDX caused immediate weight loss, followed by a rebound, and subsequent weight loss. In females, postpubertal OVX caused significant weight gain compared to sham, but prepubertal OVX did not significantly affect weight. These findings indicate that age at surgery determines the response to OVX in mice, consistent with one report that pubertal suppression of ovarian hormones does not affect weight gain in rats (Hodgson et al., 2020) but inconsistent with other reports that prepubertal OVX in rats increased weight compared to sham (Clark and Tarttelin, 1982; Freudenberger and Billeter, 1935; Freudenberger and Howard, 1937; Grant et al., 2021).

While we did not find a significant effect of prepubertal OVX on weight gain or body composition, we observed effects on food intake when operant responding was required . Both pre- and postpubertal OVX females poked less than sham females under FR1 and PR conditions. Interestingly, the timing of OVX affected FR1 performance. Prepubertal OVX females achieved a higher accuracy whereas accuracy was lower in postpubertal OVX females compared to sham. In the PR task, GDX animals across sex and age at surgery expended less effort, evidenced by significantly lower max breakpoints and reduced ramping of operant responding in PR vs. FR1. The neural circuits that regulate food motivation are believed to largely overlap with those that control drug-seeking behaviors (Ferrario et al., 2016). Given the link between striatal dopamine and effort/cost-sensitivity (Mourra et al., 2020; Salamone and Correa, 2012; Zhang et al., 2003) and gonadal hormones and dopamine signaling (Jardí et al., 2018b; Purves-Tyson et al., 2014, 2012; Yoest et al., 2018), an important future direction will be to determine whether changes in dopamine system function contribute to the reduced motivation for food we observed in GDX mice.

### Limitations of the study

In the current study, we examined feeding behavior starting 35 dps, although our current data and other studies indicate that these measures are most dynamic within ~21 dps (Blaustein and Wade, 1976; MacLean et al., 2010). Therefore, we are unable to directly relate food intake to some of the rapid changes that we observed post-surgery, for instance, weight gain following adult OVX. However, a study that compared OVX effects in adult mice vs. rats found that OVX mice gained weight in the first 21 dps in the absence of increased food intake, whereas increased food intake accompanied weight gain in OVX rats during this same period (Witte et al., 2010). At 35 dps we detected that adult OVX mice tended to consume less food relative to their intact counterparts, adding to the existing data suggesting that OVX-induced weight gain is not sustained by elevated food intake (Chen et al., 2015; McElroy and Wade, 1987; Mook et al., 1972). One study that tracked food intake in OVX mice immediately following surgery did not detect hyperphagia, and similar to our findings, observed decreased food intake in OVX mice at 8 weeks post-surgery despite higher BW (Rogers et al., 2009). The primary reason that we did not track food intake continuously after surgery is that we did not want to single house the P25 surgery group during the adolescent period when social isolation may be particularly stressful (Walker et al., 2019). The secondary reason was the practical consideration of having sufficient FED3 devices to maintain experimental throughput. In future experiments, we can either co-house same treatment siblings and compare food intake across cages or utilize methods to tag individual mice and integrate with FED3 data (Ali and Kravitz, 2018; Habedank et al., 2022). For our PR task data, we compared behavior under low-cost (FR1) vs. high-cost (PR) conditions to coarsely assess how cost influences the elasticity of food demand and effort. However, to model exponential food demand and estimate the elasticity coefficient, in the future we will need to test individual mice under multiple PR increments (PR2, PR4 etc.).

Next, we did not collect data regarding energy expenditure which will impact physiological endpoints we measured such as weight or grams of food consumed per BW. Other studies have shown that adult OVX can induce weight gain preceding or in the absence of increased food intake (Mueller and Hsiao, 1980; Richard et al., 2017; Rogers et al., 2009; Roy and Wade, 1977; Vieira Potter et al., 2012; Witte et al., 2010), pointing to changes in metabolic rate, particularly in mice (Witte et al., 2010). Physical activity is regulated by ovarian hormones in females (Berchtold et al., 2001; Isken et al., 2008; Ogawa et al., 2003; Rogers et al., 2009), and recent studies have demonstrated that estradiol’s actions via ERα on ventral medial hypothalamic (VMH) circuits promote locomotion (Correa et al., 2015; Krause et al., 2021; Musatov et al., 2007). Changes in thermoregulation, which we did not measure, are also likely to contribute to differences in energy expenditure (Dacks and Rance, 2010; Grant et al., 2021; Yochim and Spencer, 1976). Together, these data suggest that decreased energy expenditure in OVX at P60 mice are likely a major driver of the positive energy balance we observed. In males, adult ORX also causes clear reductions in locomotor activity (Butler et al., 2012; Daan et al., 1975; Dubois et al., 2016; Hoskins, 1925; Ibebunjo et al., 2011), again indicating that gonadal influences over energy expenditure likely mediated some of the observed effects. In future work we plan to incorporate home-cage measurements of activity as well as indirect calorimetry to assess changes to energy expenditure.

While our results indicate that the timing of GDX relative to puberty is important for determining the effects on energy balance, we did not perform the hormone replacement studies necessary to determine whether gonadal hormones have organizational effects during peripuberty. Finally, we did not track estrous cycle in our adult sham females during feeding experiments. However, unlike rats, the evidence that female mice exhibit estrous-cycle dependent changes in food intake is mixed (Petersen, 1976; Smarr et al., 2019; Witte et al., 2010). Future studies in our lab may carefully assess the relationship between estrous cycle and *ad libitum* vs. operant responding for food in mice.

### Summary

The major takeaway of this study is that GDX interacts with both sex and age at surgery to determine its effect on weight and body composition. On the other hand, GDX decreases food intake and food motivation regardless of sex or timing of surgery relative to puberty. In adult mice, we replicated previous findings that GDX decreases food intake and increases adiposity in males and females despite having opposing effects on weight. Our findings from prepubertal GDX mice indicate that ORX decreased weight gain and changed body composition within a short period following surgery (2.5 weeks), whereas OVX had no apparent effects on weight or body composition during adolescence. This is in striking contrast to the rapid weight gain observed when OVX was performed in adulthood. Our findings in the FR1 and PR tasks suggest that GDX decreased motivation or willingness to expend effort for food, independent of the effects it had on BW. Future studies are needed to investigate the mechanisms underlying sex-dependent effects of prepubertal GDX and address whether peripuberty may be an organizational window for energy balance regulation.

## CRediT author statement

**Klappenbach CM**: methodology, investigation, software, formal analysis, writing – original draft, and visualization **Wang Q**: investigation **Jensen A**: investigation and formal analysis **Glodosky NC**: formal analysis **Delevich K**: conceptualization, writing – review & editing, funding acquisition, and supervision.

## Acknowledgements

We thank Dr. Lex Kravitz and the FedForum for technical assistance. We thank Megan McGraw, Dr. Sara Westbrook, and other members of the Delevich lab for discussion and feedback.

