## Supplemental material for "Sex and timing of gonadectomy relative to puberty interact to influence weight, body composition, and feeding behaviors in mice"

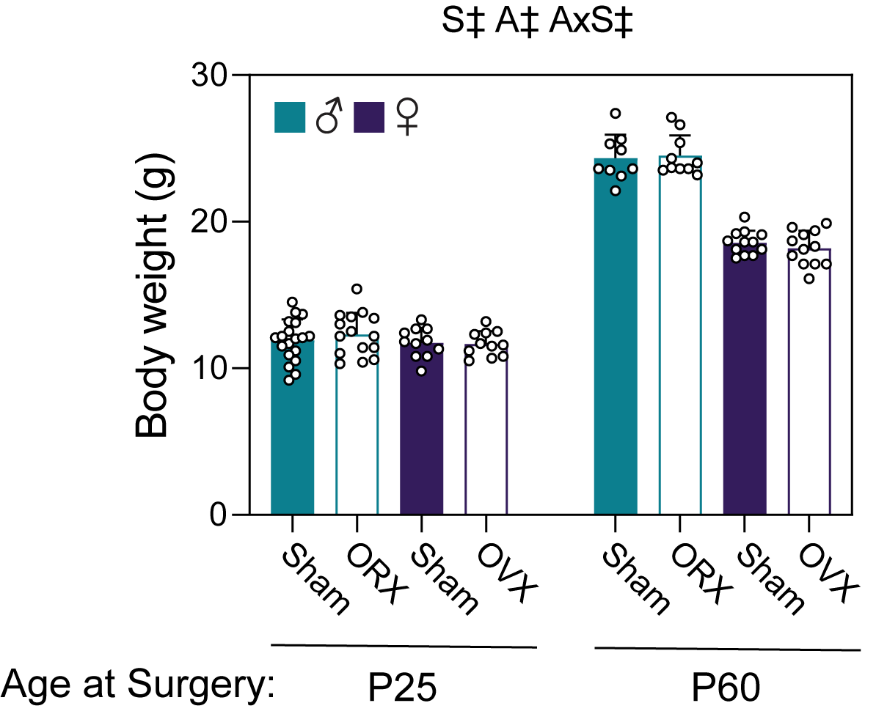

**Supplemental Figure 1**: BW was taken on the surgery day to ensure there were no preexisting differences between sham and GDX animals. These results also show that sex differences in BW emerge post puberty. Statistics above panel represent a three-way ANOVA between gonadal status (G), sex (S), and age of surgery (A) where ‡ = *p* < .0001, and bars represent mean ± SEM, n = 9-19 mice / group.

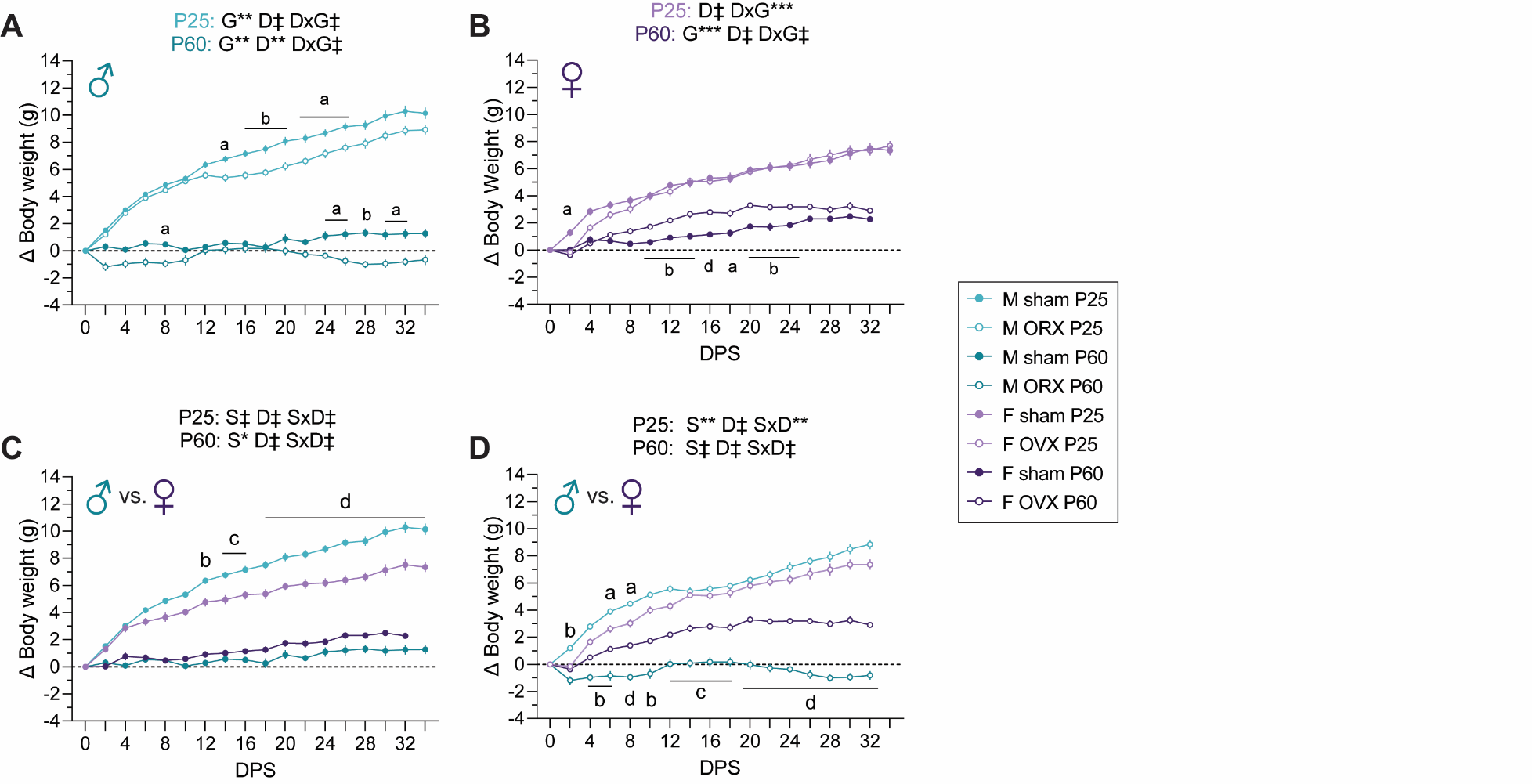

**Supplemental Figure 2:** BW gain following pre-pubertal (P25) or post-pubertal (P60) gonadectomy or sham surgery. **A:** Effect of sham vs ORX surgery in males. **B:** Effect of sham vs OVX surgery in females. **C:** Comparison of weight gain in sham animals. **D:** Comparison of weight gain in GDX animals. Statistics above the figure panels represent two two-way ANOVAs within surgery age. Significant main effects and interactions of gonadal status (G), sex (S), and days post-surgery (D) are indicated by * *p* < .05, ** *p* < .01, *** *p <* .001, and ‡ *p* < .0001. Lowercase letters indicate significant *post hoc* comparisons within age and days post-surgery using Sidak’s multiple comparison test where a–d indicate *p* < .05, *p* < .01, *p* < .001, and *p* < .0001, respectively. Points represent mean ± SEM, n = 9-17 mice / group.

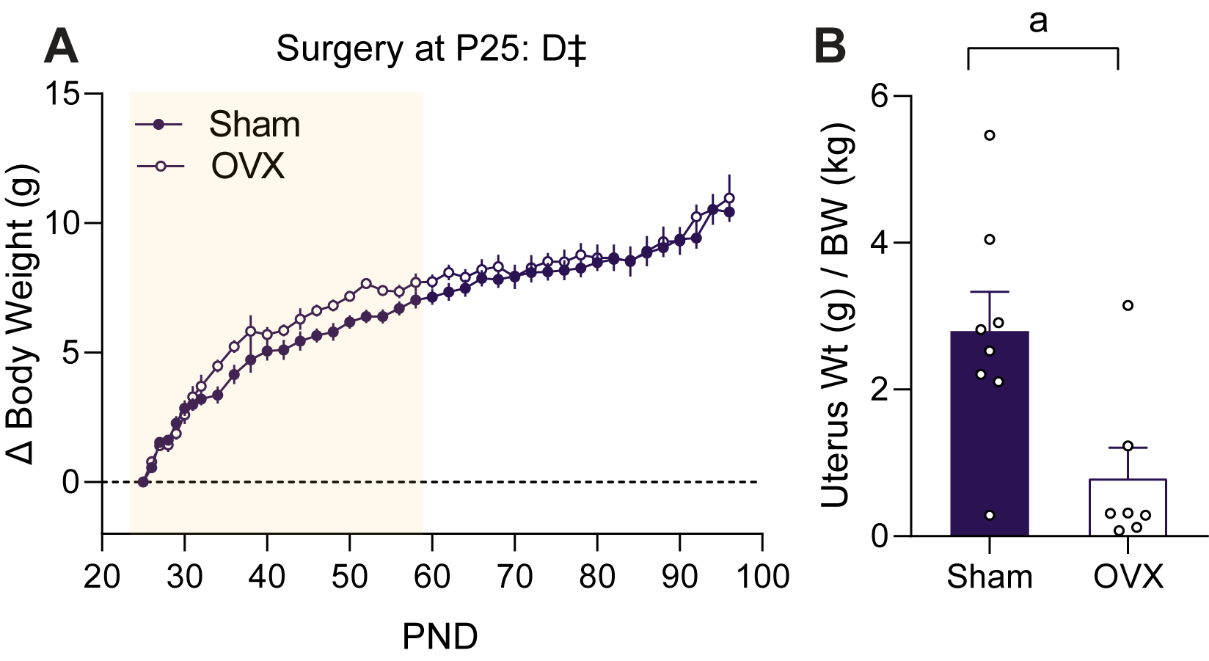

**Supplemental Figure 3:** In a separate cohort of mice, sham or OVX surgery was performed pre-pubertally (P25) and BW was measured for 65 DPS. **A.** BW gain following surgery. Points represent mean ± SEM**.** Statistics above panel represent two-way ANOVA between surgery and DPS (D) where ‡ = *p* < .0001. n = 7-10 mice / group. **B:** Weight in g of uterus normalized to BW in kg at the end of BW measurements. Bars represent mean ± SEM**.** n = 7-8 mice / group. An unpaired t test between sham and OVX mice was performed and a represents *p* < .05. One OVX mouse was removed from both panels as a result of ROUT outlier analysis in normalized uterus weight (Q = 1%).

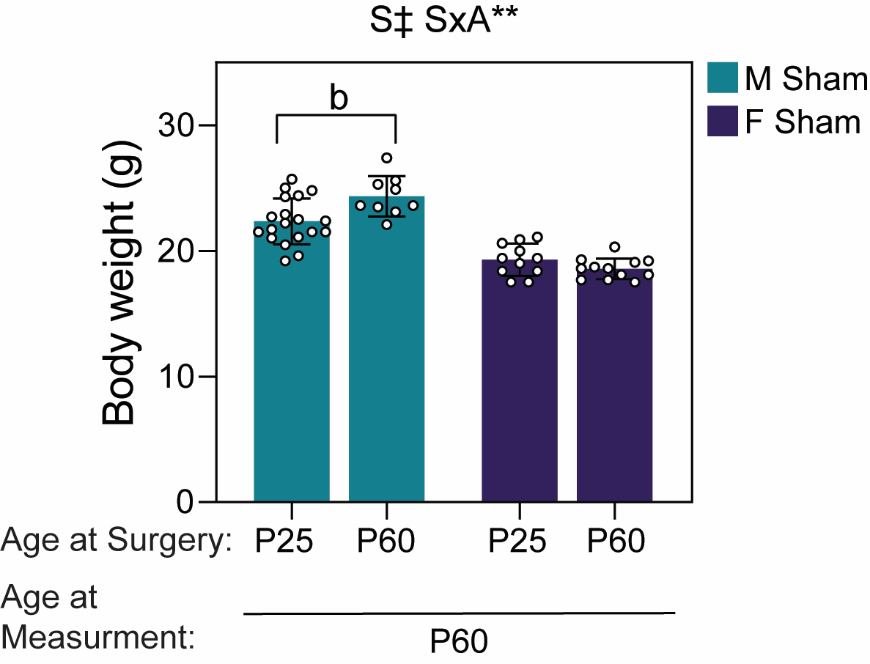

**Supplemental Figure 4:** Sham surgery at P25 decreases P60 weight in male but not female mice. Comparison of sham surgery groups indicated a significant interaction of sex and age at surgery with P25 sham males weighing less at P60 than the P60 sham group (pre-surgical weight). Statistics above panel represent significant main effect of sex (S) and a significant interaction between sex (S) and age at surgery (A) (two-way ANOVA). ** *p* < .01, ‡ *p* < .0001, and b indicates *p* < .01 from *post hoc* comparison using Sidak’s multiple comparison test. Bars represent mean ± SEM. n = 9-19 mice / group.

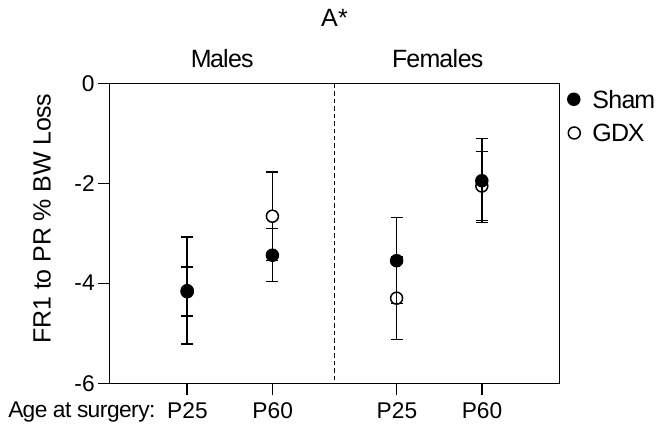

**Supplemental Figure 5:** Percent weight loss in mice switching from FR1 to PR. Calculated from weight immediately prior to starting PR and weight after 24 hours of PR. n = 9-18 mice / group. Statistics above panel represents significant main effect from a three-way ANOVA between sex, age of surgery (A), and gonadal status.
